# Action intentions reactivate representations of task-relevant cognitive cues

**DOI:** 10.1101/2024.11.29.626095

**Authors:** Nina Lee, Lin Lawrence Guo, Adrian Nestor, Matthias Niemeier

**Affiliations:** Department of Psychology at Scarborough University of Toronto Scarborough, ON, Canada, M1C1A4; Centre for Vision Research York University Toronto, ON, Canada, M4N3M6

## Abstract

Recent research shows that the intention to act on an object alters its neural representation in ways as afforded by underlying sensorimotor processes. For example, the intention to grasp and pick up an object results in representations of the object’s weight. But these representations become grasp-specific only immediately before object lift if weight information is relayed through object material. This feature triggers earlier representations regardless of intention probably because material-weight contingencies are overlearned. By contrast, recently learned weight cues should be recalled deliberately during grasp planning resulting in early grasp-specific representations. Here, we examined how action intentions affect the representation of newly acquired colour-weight contingencies. We recorded electroencephalography while human participants grasped or reached for objects that varied in shape and density as indicated by their colour. Multivariate analyses revealed a grasp-specific reactivation of colour during planning that was mirrored in beta band. This suggests that task-relevancy influences the representation of colour such that previously encoded colour-weight contingencies may be reactivated as required for grasping, mediated top-down via working memory. Grasp-specific representations of shape and colour were also present in theta band, perhaps reflecting attentional activity. These results provide novel insights into the interplay between cognition and motor planning processes.

**Significance Statement:** Recent research shows that the object feature weight, conveyed via highly overlearned material-weight contingencies, is more prominently represented within electrophysiological signals when people intend to grasp an object to lift it vs. reach for it. However, such differences occur late, immediately before object lift. Our study is the first to show that newly learned colour-weight contingencies yield early task-specific representations during action planning; we find that colour/weight representations, forming 140-210 ms after object onset, reactivate around 270 ms during grasp, but not reach planning, with the same pattern arising for beta band. These data suggest that the brain deliberately processes weight information depending on intention, arguably involving working memory representations. Our results highlight the interplay of cognition and motor planning.

## Introduction

“I suppose it is tempting, if the only tool you have is a hammer, to treat everything as if it were a nail.” While Maslow’s (1966) statement referred to the biasing influence of task sets on cognitions, task sets also bias sensory and sensorimotor processes. The way we see the world depends on how we intend to act (e.g., Bekkering & Neggers, 2002; Craighero et al.,1999; Fagioli et al., 2007; Shin et al., 2010). More specifically, action intentions such as grasping have been associated with altered spatial attention (Brouwer et al., 2009; Schiegg et al., 2003; Baldauf & Deubel, 2009; Baldauf et al., 2006) and action-specific activation of early visual areas (Gallivan et al., 2019), arguably through sensorimotor feedback signals from parietal cortex (e.g., Andersen & Cui, 2009; Cui, 2014).

Consistent with feedback signals activating early cortex, multivariate electroencephalography (EEG) studies have recently shown that human participants form early neural representations of the shape of an object ∼100-250 ms after it appears and that these representations re-occur after ∼250 ms (Guo et al., 2019), but only when the participants intend to grasp the object, not when they plan to simply reach for it (Lee et al., 2024). These reactivation patterns of shape representations suggest that grasp intentions trigger a detailed visual analysis of grasp-relevant object features to extract information about how to guide the grasping fingers to the object during movement planning.

Around the same time of planning, grasp-relevant information about the object’s weight has been found to form neural representations, too (Lee et al. 2024). However, these representations were largely the same regardless of whether participants intended to grasp or reach for objects. Only during movement execution, shortly before participants lifted an object, grasp-specific representations were observed (Lee et al., 2024; also see Klein et al., 2023). This suggests that grasp-specific computations of object weight are confined to the respective grasping phase when the weight of an object becomes relevant. However, the onset of grasp-specific computations might depend on what kind of weight cues are available.

Stable weight cues probably are heavily overlearned because they are frequently and consistently encountered in daily life. For example, Lee and colleagues (2024) used objects that were made of wood or steel (also see Gallivan et al., 2014). These materials are so common in daily life that weight information might be automatically generated as soon as the material becomes visible – regardless of whether weight information is relevant for an intended action. By contrast, arbitrary cue-weight contingencies (e.g., Ameli et al., 2008) might not be recalled automatically because they are not overlearned. For example, arbitrary colour-weight contingencies, learned within the short time span of an experiment (e.g., Li et al., 2009), should be recalled only when they are relevant for the intended action. Furthermore, given that these arbitrary contingencies require cognitive resources (Baugh et al., 2016; Li et al., 2009) the respective cognitive processes, arguably involving representations in working memory (Gallivan et al., 2014; Guillery et al., 2017; Li et al., 2009; Zhang et al., 2023), should arise in an action-specific manner.

It follows that specifically when people intend to grasp an object, transient colour weight cues should trigger memory queries about the contingency, probably as soon as an object becomes visible and the resulting grasp-specific representations of colour might be modulated top-down by working memory (e.g., Lundqvist et al., 2011; Richter et al., 2018; for review: Miller et al., 2018). Thus, we expected that specifically for grasping EEG signals should carry information about object colour, probably in the form of reactivation patterns as previously observed for shape information (Guo et al., 2021; Lee et al., 2024).

Indeed, this information might be contained in beta band oscillations which have previously been implicated in the top-down modulation of sensory processing and working memory maintenance (Fiebelkorn & Kastner, 2019; Lundqvist et al., 2011; Richter et al., 2018), as well as the reactivation of task-relevant working memory content (Spitzer et al., 2014; Spitzer & Haegens, 2017; Wimmer et al., 2016). Theta band might be another promising candidate given its association not only with working memory processes (Hsieh & Ranganath, 2014; Sauseng et al., 2010; Tesche & Karhu, 2000), but also goal-directed and feature attention (Fellrath et al., 2016; Harris et al., 2017; Liebe et al., 2012).

To test these predictions, in the current study we presented participants with 3D-printed objects of different colours signaling different densities and, thus, weight and asked them to grasp or to merely reach for them. Our aim was to this way elucidate the interactions of the sensorimotor processes underlying grasp actions and cognitions so as to explore the immense adaptability of behaviour under the guidance of human thought.

## Materials & Methods

### Participants

Twenty undergraduate and graduate students (11 females; mean age: 22.6) from the University of Toronto Scarborough gave their written and informed consent to participate in the experiment and were compensated $15/h for their time. All participants had normal or corrected-to-normal vision and were right-handed (Oldfield, 1971). All procedures were in accordance with the Human Participants Review Sub-Committee of the University of Toronto (approval ID: 37329 for the neural and cognitive mechanisms underlying predictions and attention) and with the Declaration of Helsinki involving human participants.

### Stimuli

We introduced participants to four objects that differed in shape and colour. Objects were blue and red 3D-printed “pillows” and “flowers” that were 2 cm in depth (Fig. 1C). Pillows had concave (72 deg. segments of circles with a radius of 7.5 cm, i.e., curvature = 13.3/m), and flowers had convex sides (180 deg. segments of circles with a radius of 1.5 cm, curvature = 66.7/m). These shapes were chosen to have visually distinct surface curvatures, while at the same time, all objects measured 6 cm across opposing edges, thus affording identical grip sizes. Critically, for a given participant, object sets (i.e. one pillow and one flower) of one colour (i.e. red or blue, counterbalanced across participants) were hollow and thus 48% lighter compared to shapes of the opposing colour (heavy pillow: 120 g, light pillow 62 g; heavy flower: 63 g; light flower: 33 g). In this way, colour was a reliable indicator of object density and weight. Thus, colour, just like shape, was a visual feature that was relevant when we asked participants to grasp objects with their thumb and index finger, but not in the control condition where participants reached and touched the centre of the objects with the knuckle of their index finger while making a fist (subsequently called “knuckling” in text).

**Figure 1.**
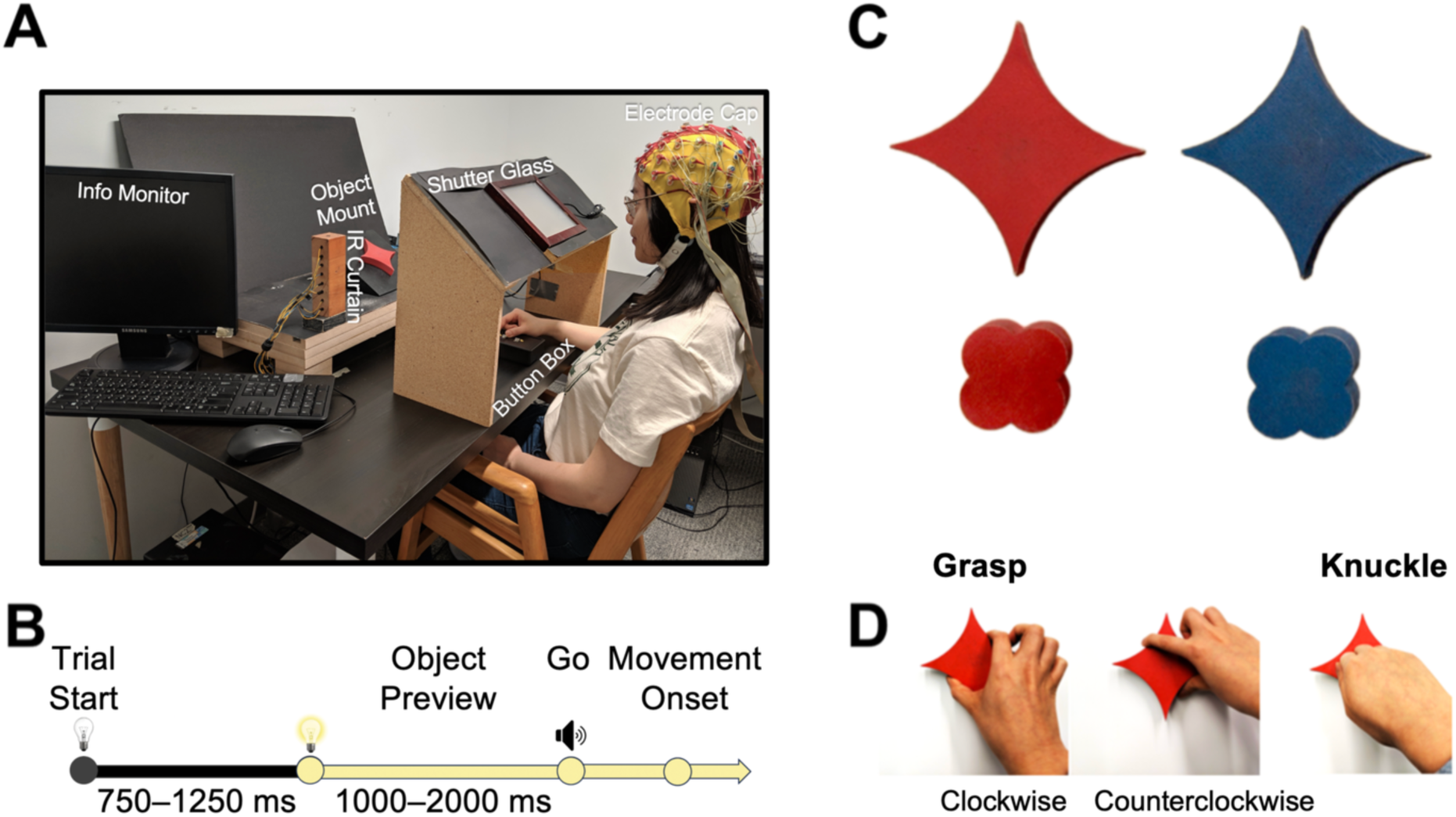
Experimental methods. ***A,*** Experimental setup. ***B***, Timeline of a trial. ***C***, Objects used in the experiment. ***D***, Action conditions.

### Procedure

Prior to the start of the experiment, participants were given a chance to lift each of the four objects. The experimenter explicitly drew attention to the fact the objects of one colour were heavier than those of the other colour. During each session, then participants sat in a dark room. The experimenter, seated on their left (Fig. 1A), prepared each trial; they viewed instructions on a monitor and turned on a pair of LEDs to mount objects on a platform in an otherwise dark grasp space. The platform was slanted toward the participant’s line of sight, and a square-shaped hole in its centre ensured that the objects, with a square peg attached to their back, were always placed in the same position and orientation, 43 cm away from the participant immediately behind a curtain of infrared beams (created by two 15 cm tall pillars, 40 cm apart). Relative to the bottom of object, the infrared curtain was 6 cm away. This pre-trial set-up could not be monitored by the participant because they wore earplugs and had their view blocked by an opaque shutter glass screen (Smart Glass Technology). The participants then placed their right index finger and thumb on a button box, each covering a beam of infrared light to sense the presence of their hand. Next, the experimenter pressed a key that turned off the LEDs and, in darkness, set the shutter glass screen to transparent. Seven-hundred and fifty to 1250 ms later, the LED lights turned on, thus illuminating the object for the participant to see for a “Preview” duration of 1-2 seconds before an auditory “Go” tone (loud enough to hear through the earplugs) signalled the participant to make their movement. As participants lifted their hand off the button box, this was recorded as movement onset, and the hand intersecting the infrared curtain was recorded as movement end. The kind of movement to be made (clockwise or counter-clockwise precision grasps and lifts, or knuckles with a comfortable horizontal wrist orientation) was announced at the beginning of each block of trials (Fig. 1D) and monitored by the experimenter (they marked invalid trials when participants moved incorrectly or dropped objects; n.b., we included the variable Orientation merely to be able to compare to previous studies, Guo et al., 2019; Guo et al., 2021; Lee et al., 2024). There were 72 trials (2 shapes, 2 colours x 18 repetitions in random order) in one block, and there were 18 blocks in total (six for each movement type: clockwise grasps, counter-clockwise grasps, and knuckles) administered in random order and in 2 sessions of about 3 hours each over two different days with breaks being provided between blocks as needed.

### EEG Acquisition and Preprocessing

We recorded EEG data using a 64-electrode BioSemi ActiveTwo recording system (BioSemi B.V., Amsterdam, Netherlands), digitized at a rate of 512 Hz with 24-bit A/D conversion. The electrodes were arranged according to the International 10/20 System, and the electrode offset was kept below 40 mV.

EEG preprocessing was performed offline in MATLAB using the EEGLAB (Delorme and Makeig, 2004) and ERPLAB toolboxes (Lopez-Calderon and Luck, 2014). Signals were bandpass filtered (noncausal Butterworth impulse response function, 12 dB/oct roll-off) with half-amplitude cut-offs at 0.1 and 40 Hz. Next, noisy electrodes which correlated <0.6 with nearby electrodes were interpolated (mean of 1.6 electrodes per subject), and all electrodes were re-referenced to the average of all electrodes. Independent component analysis (ICA) was then performed on continuous blocks for each participant to identify and remove blink, and eye movements components (Jung et al., 2000; Chaumon et al., 2015; Drisdelle et al., 2017). The ICA-corrected data were segmented relative to the onset of Preview (-100 to 1200 ms). In addition, invalid trials and epochs containing abnormal reaction times (less than 100 ms or greater than 1000 ms) were removed. As a result, an average of 4.1% of trials from each subject were removed from further analysis.

### Pattern Classification of ERP Signals Across Time

We averaged epochs into ERP traces to increase signal-to-noise ratio (SNR) of spatiotemporal patterns (Grootswagers, Wardle, & Carlson, 2016). Specifically, we pooled together blocks of the same action (e.g. clockwise grasping). Up to 18 epochs within a given action block that corresponded to the same object (e.g. red pillow) were averaged together, resulting in 6 separate ERP traces per condition (e.g., clockwise grasping of red pillows) for Preview. We then performed multivariate noise normalization by calculating a covariance matrix for all time points of an epoch separately for each condition, then taking the mean of these covariance matrices across time points and conditions (Guggenmos, Sterzer, & Cichy, 2018). Next, we *z*-scored traces across time and electrodes, and outliers were thresholded at ±3 SD from the mean (for similar approaches see Nemrodov et al., 2017; 2018). We opted for this winsorization approach to reduce the impact of outliers on support vector maching (SVM) pattern classification while maintaining the same number of features.

Further, we divided ERP traces into temporal windows with 5 consecutive bins (5 bins * 1.95 ms ≈ 10 ms) to increase the robustness of SVM analyses. For each time bin, data from all 64 electrodes were concatenated to create 320 features. In this way, we could perform time-resolved pattern classification across time through these bins.

For all decoding analyses we first performed classification for each participant separately and then determined the mean performance across participants. Using SVM we conducted pairwise discrimination of Shape, Colour, hand Orientation (on grasping trials only), and Action with a leave-one-out cross-validation approach (i.e. through iterations, all data were trained and tested on). More specifically, for Action, 35 of 36 pairs of observations were used in training, while 1 pair was used for testing. Due to the imbalanced nature of the number of grasping trials versus knuckling trials (i.e. there were twice as many grasping compared to knuckling trials), we set the cost parameter to be double the weight of the majority class (c = 2 vs. c = 1; Batuwita & Palade, 2013) to avoid the classifier preferentially skewing towards the majority class. Further, we ensured the testing sample contained an equal number of minority (knuckling) and majority (grasping) instances. For the other independent variables, we separately analyzed grasping and knuckling trials. For Shape, Colour and Orientation, in the case of grasping: 23 of 24 pairs of observations were used for training and 1 pair was used for testing. For Shape and Colour classification during knuckling: 11 of 12 pairs of observations were used for training and 1 pair was used for testing (there was not an equivalent Orientation condition for knuckling). For all Action and Orientation analyses, we created ERPs from randomly drawn blocks to account for blocked effects (due to action and hand orientation being held constant in a block, whereas objects varied by trial).

Further, to test for potential integrated representation of task features, we performed pairwise discrimination for Shape ∩ Colour, separately for grasping and knuckling similar to Lee et al. (2024). More specifically, we trained and tested the SVMs with the following logic to reduce the possibility that classifiers relied only on one feature (i.e., shape or colour) to classify: for the integrated representation of interest (e.g. “red pillow”) we included training and testing data that shared only the shape feature (e.g. “blue pillow”), only the colour feature (e.g. “red flower”), or both features (e.g. “red pillow”). We treated this as an instance of imbalanced classes (see Action classification above) such that data containing both features would be considered the minority class, whereas data containing one feature would be considered the majority class. This resulted in a cross-validation setup of 33 observations used for training and 2 for testing for grasping and 15 observations for training and 2 for testing for knuckling. We trained four separate classifiers using this logic (i.e. the exhaustive combination of shape and colour) and averaged their performance to obtain an estimate of Shape ∩ Colour representations.

Next, we tested this averaged performance against an optimized additive effect between Shape and Colour to determine whether these classifiers truly reflected integrated representations. To this end, we first calculated how a single-feature classifier would perform in our test:

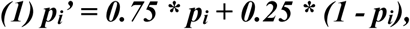

where ***p_i_*** is the accuracy of some single-feature classifier ***i*** and ***p_i_’*** is its performance in our test (e.g., a shape classifier that attains an accuracy of p_shape_=0.7 would classify a minority object “red pillow” and a majority object “red flower” as belonging to class 1 and 2, respectively, with p=0.7 each, and it would classify the second majority object “blue pillow” with p=1-0.7 as belonging to class 2. This would result in correct classification with p_shape_’ = 0.5 * 0.7 + 0.25 * 0.7 + 0.25 * (1-0.7) = 0.75 * p_shape_ + 0.25 * (1-p_shape_) = 0.6).

To then obtain an estimate of how two single-feature classifiers would perform optimally together, we modelled the response of each single-feature classifier as a continuous value with a Gaussian distribution where ***µ*** is the mean, ***σ_i_*** is the standard deviation, and ***p_i_’*** is the area under the Gaussian up to a decision boundary ***x***. The p_i_’ value can then be written as a cumulative Gaussian:

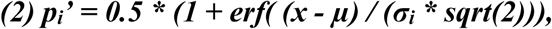

where ***erf*** is the error function and ***sqrt*** is the square-root. Solving equation (2) for σ_i_ for both single-feature classifiers (and with arbitrary values for µ and x), allows us to use maximum likelihood estimation to determine the optimally combined standard deviation for both single-feature classifiers together:

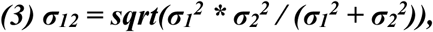

where ***σ_1_*** and ***σ_2_*** are the standard deviations for the two single-feature classifiers, respectively, and ***σ_12_*** is the standard deviation of the optimally combined classifier. Inserting ***σ_12_*** into equation (2) gives us the optimal performance of both classifiers together (one exception: if one or two classifiers performed at guessing rate or worse, optimal performance p_12_ was set to the maximum of p_1_, p_2_, or 0.5). This way, the accuracy of the integrated classifier was tested relative to p_12_.

### Cross-Temporal Generalization of ERPs

We used cross-temporal generalization of ERPs to investigate whether classifiers trained on a one time point can decode data from the same or different time points. Thus, we trained and tested classifiers on all combinations of times from the first 500 ms of Preview (Fig. 3). We shortened the time window as analyses beyond 500 ms did not yield significant results. To further examine the differences and similarities of how actions modulate representations, we conducted separate analyses: training and testing on grasping, training and testing on knuckling. If there were similarities in the representation of two selected time windows, the classifier should generalize between them. More specifically, there are three possible patterns of classifier performance we expect to see: 1) Chained representations. Represented by a classification pattern along the diagonal of the graph, the classifier should perform above chance only when trained and tested on the same time points. Transient neural representations that activate sequentially would likely result in this pattern. When seen in isolation, this would suggest the feature does not share representations across time and would be inconsistent with our hypothesis of encoded feature information that is used at later times. 2) Reactivated representations. Represented by significant classification along the diagonal as well as in approximately symmetrical “arms” above and below the diagonal. This means that the classifier trained at an earlier time performs above chance for later time points, reflecting that later neural representations are reactivated versions of earlier ones. We hypothesized to see this pattern of representation for Grasping and not Knuckling for Shape and Colour, reflecting the requirement of earlier encoded feature information to successfully carry out the grasp. 3) Sustained representations. Represented by a wide coverage of areas above and below the diagonal, the classifier trained at an earlier time will perform above chance when tested for times immediately following the training time. Such a pattern of generalization would implicate a stable neural code or process over time. Although it would not necessary be inconsistent with our hypothesis, it would be unexpected based on past work to find this pattern for visual features (Guo et al., 2019; 2021; 2024; Lee et al., 2024). Instead, sustained representations have been found for highly robust features like action or grasp/reach orientation.

### Pattern Classification of Time Frequency Data

We performed classification of time frequency transformed data by first bandpass filtering (delta: 1-3 Hz; theta: 3-8 Hz; alpha: 8-12 Hz; beta: 15-25 Hz; gamma: 30-40 Hz) epochs using two-way least-squares finite impulse response filtering. We applied Hilbert transform on narrowband data to obtain the discrete-time analytical signal and squared complex values to obtain the power. This band-separated power was grouped, normalized and averaged in the same way as ERP traces for spatiotemporal EEG classification (see “Pattern Classification of ERP Signals Across Time” and “Cross-Temporal Generalization of ERPs”).

### Electrode Informativeness

We determined the informativeness of electrodes for the classification of ERP effects (Shape, Colour, Shape ∩ Colour, Orientation, and Action for data aligned to Preview) by conducting a searchlight analysis across electrodes. For each electrode we defined a 50 mm radius neighbourhood and performed classification on spatiotemporal features obtained from each electrode neighbourhood across 100 ms time bins. For Shape ∩ Colour, the values at each electrode were subtracted from the respective values obtained from the optimal integration of both classifiers together at each time bin and at each electrode (section “Pattern Classification of ERP Signals Across Time”).

### Significance Testing

For behavioural data we determined statistical differences (reaction and movement time) using three-way repeated measures ANOVAs (Action x Shape x Colour), adjusting for sphericity violations using Greenhouse-Geisser (GHG) corrections. We assigned Action as a three-level factor (i.e. clockwise grasping, counter-clockwise grasping and knuckling) for completeness, but we did not find any difference between the two grasping conditions and subsequently did not treat these as separate groups for classification analyses. For post hoc analyses, we used repeated measures t-tests with Bonferroni correction for multiple comparisons.

For all analyses conducted on EEG data we assessed statistical significance using a non-parametric, cluster-based approach to determine clusters of timepoints (or electrodes for searchlight analyses) where there were group-level significant effects (Nichols and Holmes, 2002). For example, in the case of time-resolved analyses, we defined clusters as consecutive time points that exceeded the 95^th^ percentile of the distribution of t-values at each time point attained using sign-permutation tests computed 10000 times, equivalent to p<0.05, one-tailed. Further, significant clusters were determined by cluster sizes equal to or exceeding the 95^th^ percentile of maximum cluster sizes across all permutations (equivalent to p<0.05, one-tailed). A similar approach was taken for searchlight analyses carried out across electrodes, and cluster-based correction was performed on each 100 ms time window, separately on spatial clusters of electrodes.

In the case of temporal generalization analyses, we tested for significant differences between grasping and knuckling for the three expected types of representations (i.e., chained, reactivated, and sustained representations), as well as a baseline (the first 50 ms, when representations are not yet expected in visual cortex) during Preview (Fig. 3). We obtained these regions of interest (ROIs) from independent criteria (see Lee et al., 2024, Significance Testing for cross-temporal generalization), which will henceforth be referred to as “diagonal” ROIs (indicative of chains of multiple neural generators of the representations), “arms”-shaped ROIs (indicative of generators that reactivate at a later time), and “triangular” ROIs (indicative of generators that are active for a longer, sustained time; King & Dehaene, 2014).

For each of these ROIs we obtained corresponding classification accuracy values from grasping and knuckling trials separately and then calculated a difference score of the average grasping vs. knuckling accuracies at the respective ROIs. We then determined the one-tailed 95% confidence intervals of these differences (because a priori we expected representations during grasping to be more pronounced than representations, if any, during knuckling) using non-parametric bootstrapping with 10000 iterations. For the diagonal, arms-shaped, or triangular ROI differences to be significant, these values had to be more positive than the difference for the baseline ROI with no overlap in confidence intervals to ensure that classification was better than based on unspecific differences between experimental conditions. We completed this significance testing process for all grasping vs. knuckling comparisons of main effects and integrated representations. A significant difference within the diagonal ROI would suggest a stronger chained representation. A significant difference within the “arms”-shaped ROI (but not the triangular ROI) would suggest a stronger reactivated representation for grasping than knuckling. Furthermore, a significant difference within the triangular ROI (with or without differences in the arm-shaped ROI) would signify a stronger sustained representation for grasping compared to knuckling.

## Results

### Behavioural Results

The mean reaction time (RT; defined as the time between Go onset and movement onset) was 286 ms (SD = 72 ms), and mean movement time (MT; defined as the time between movement onset and movement end) was 244 ms (SD = 49 ms). None of the main effects or interactions of the ANOVA for RT were significant (F’s < 1.798, p’s > 0.196). However, there was a main effect of Action for MT (F (2, 38) = 22.200, p < 0.01 η_p^2 = 0.539). Pairwise comparisons with Bonferroni correction revealed that knuckling trials had slower MTs (M = 272, SD = 41 ms) compared to both clockwise grasping (M = 226 ms; SD = 41 ms) and counter-clockwise grasping (M = 235 ms; SD = 52 ms), but grasping conditions were not significantly different from one another. All other main effects and interactions were non-significant (F’s < 0.854, p’s > 0.367).

### Investigate When Features are Represented Using Time-resolved Classification of ERPs

Time-resolved Grasping Colour representations became significant from 90 – 230 ms, reaching a maximum at 160 ms (Fig. 2A, first row, dark red), whereas Knuckling Colour representations were significant much later, from 220 – 290 ms and peaking at 260 ms (Fig. 2A, first row, light red), although there were no differences between Grasping and Knuckling.

**Figure 2.**
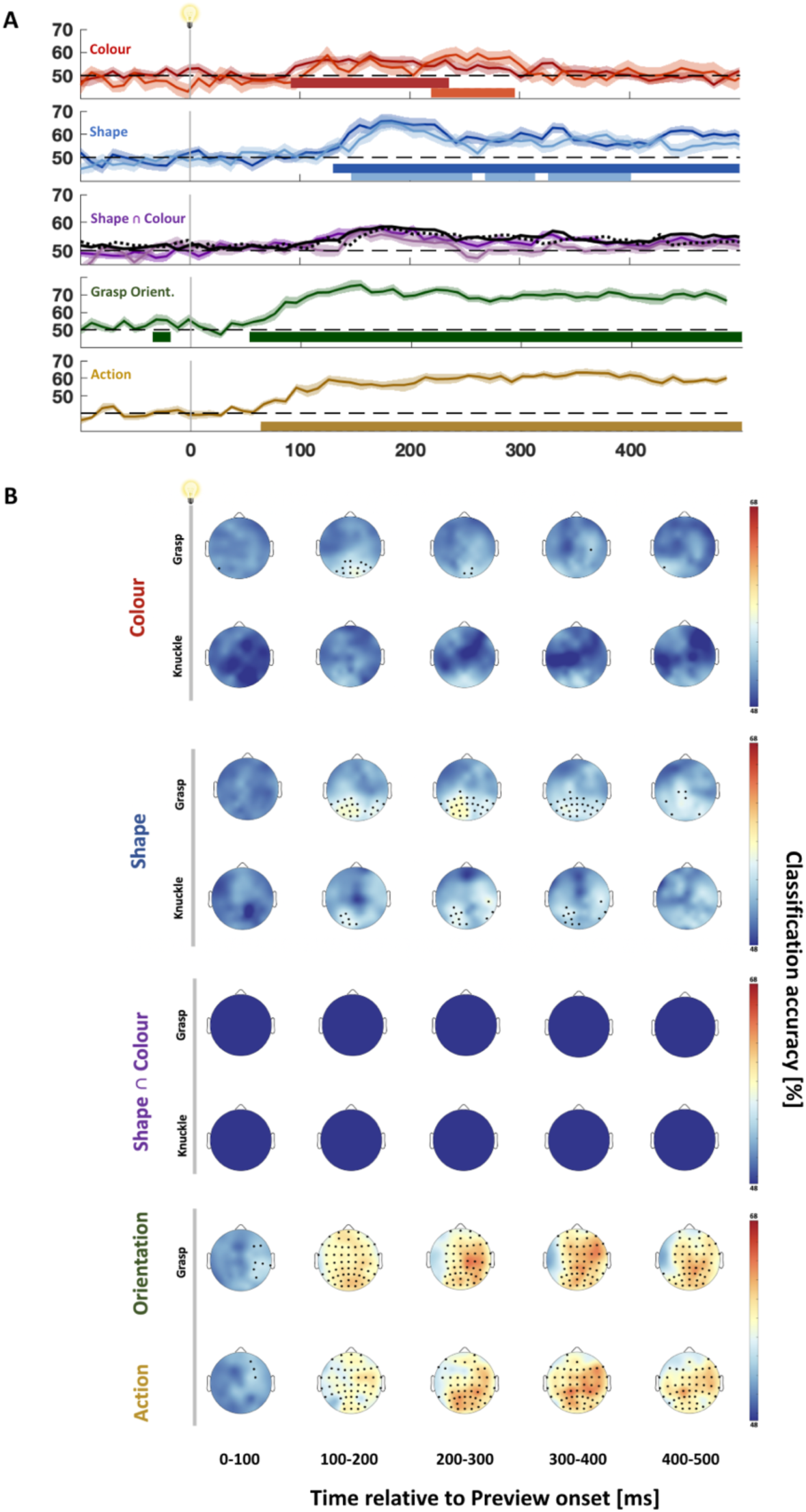
Classification accuracy of main effects and Shape ∩ Colour during Preview. **(A)** Time resolved classification of effects aligned to Preview onset. Darker curves represent classification of grasping data, lighter ones represent knuckling data. Black curves in the third graph show performance as predicted by the respective two single-feature classifiers for grasping (solid curves) and knuckling (dotted curves). Horizontal coloured bars denote significantly above chance classification (cluster-corrected t-tests, one-tailed; p < 0.05). **(B)** Electrode informativeness for respective effects aligned to Preview (two-tailed one-sample t-test; q < 0.05). No Shape ∩ Colour classifiers performed better than optimal additive models of single classifiers.

There also were no significant differences between Grasping vs. Knuckling Shapes, although both conditions were significantly different from chance. In detail, Grasping Shape representations became significant starting 130 ms after object onset, quickly rising to a peak at 180 ms and remaining significantly above chance for the rest of the Preview phase (Fig. 2A, second row, dark blue). In comparison, Knuckling Shape representations became significant slightly later than Grasping representations at 150 ms, though also reaching a peak at 180 ms. These Knuckling Shape representations were significant until 400 ms after Preview Onset (breaks at 250 – 270 ms and 310 – 330 ms; Fig. 2A, second row, light blue).

Addressing the question of whether there were integrated representations of Shape and Colour, we tested classification accuracies of Shape ∩ Colour against optimal additive models of single feature classifiers (Shape and/or Colour classifiers, Fig. 2A, third row; see “Pattern Classification of ERP Signals Across Time” for description of models). Neither Grasping Shape ∩ Colour nor Knuckling Shape ∩ Colour representations came out significant against the optimally integrated additive models, and moreover, there were no significant differences between grasping and knuckling trials.

Finally, merely to be comprehensive we also examined the two remaining experimental manipulations, Orientation and Action. We found that representations of Grasping Orientation (i.e., clockwise vs. counter-clockwise grasping; n.b., no Knuckling Orientation conditions were tested) were briefly significant before Preview Onset (-30 to -20 ms) and more substantially from 60 ms after Preview Onset, rising to a peak at 150 ms and remaining robustly significant until the end of the Preview period (comparable to Guo et al., 2019; Guo et al., 2021; Lee et al., 2024).

Classification of Action (i.e. grasping vs. knuckling) became significant at 70 ms, first rising quickly, then continuing to gradually rise and becoming sustained until the end of Preview (highest classification accuracy was at 360 ms; Fig. 2A, 5^th^ row; similar to Lee et al., 2024).

### Approximate Spatial Sources using Electrode Informativeness

To explore where information originated, we trained classifiers with a searchlight approach (individual electrode data plus their immediate vicinity). Grasping Colour representations were significant at posterior electrodes starting from 100 – 200 ms (peak electrodes: Oz, O2 and PO4 which remained significant at 200 – 300 ms), implicating visual areas. No electrodes reached significance for Knuckling Colour representations (Fig. 2B, first and second rows).

Grasping Shape information mostly originated from posterior electrodes, with peak classification lateralized to the left starting during the 100 – 200 ms interval and becoming most informative at 200 – 300 ms (peak electrodes including O1, PO3 and P3; Fig. 2B, third row), again implicating visual areas. Similarly, Knuckling Shape representations originated from posterior electrodes, with peak classification mostly on the left posterior side, though C6, TP8 and P8 also became significant between 200 – 400 ms (Fig. 2B, fourth row).

In the case of integrated Shape ∩ Colour representations, no electrodes were significant after correcting for additive models (Fig. 2B, fifth and sixth rows).

Grasp Orientation information in contrast, was widely dispersed, being most significant at posterior electrodes from 100 – 200 ms, lateralizing to the right from 200 – 300 ms (peak electrodes: C2, C4, and CP2) as well as 300 – 400 ms (peak electrodes: CP2, FC4, and FC6), and 400 – 500 ms (peak electrodes: C4, P2, and P2), suggesting a greater role of premotor and motor regions in the representation of orientation.

Action representation was also widely dispersed. From 100 – 200 ms, peak electrodes were lateralized to the right at fronto-central regions (including FC2, FC4, and FC6). Starting from 200 ms, there was greater involvement of posterior and centro-parietal electrodes (from 200 – 300 ms peak electrodes on the left side: PO3, P1 as well as right side: CP4; from 300 – 400 ms peak electrodes also included left and right sides: CP1, CP4 and F6; from 400 – 500 ms, CP1 featured most prominently), consistent with what would be expected of representations that would involve more premotor and motor regions.

### Investigate how Representations Change over Time Using Temporal Generalization of ERPs

Central to the current study and informed by our previous work (Lee et al., 2024), we expected task-dependent differences between Grasping and Knuckling to arise from temporal generalization analysis. That is, we anticipated differences between these conditions in how the representations of experimental variables changed over time. We hypothesized to see grasp-specific reactivation of visual features. Further, Lee et al. (2024) did not find a reactivation of weight during action planning when this information was relayed through overlearned material cues. Here, we expected to see evidence that weight obtained from newly associated colour cues will be represented during action planning as part of the maintenance of this weight information.

Examining Grasping Colour representations, we observed significance along the diagonal (from 110 – 170 ms along the x-axis and 110 – 160 ms along the y-axis in Fig. 3A, first graph) while no significance emerged for Knuckling Colour (Fig. 3A, second graph). Crucially, comparing differences between Grasping vs. Knuckling data for our a-priori defined ROIs revealed that Colour representations along the “arms”-shaped ROI were stronger for grasping than knuckling (Fig. 3A, third and fourth graph), suggesting that the intention to grasp an object (more so than the intention to knuckle an object) caused neural generators of colour representations to reactivate.

**Figure 3.**
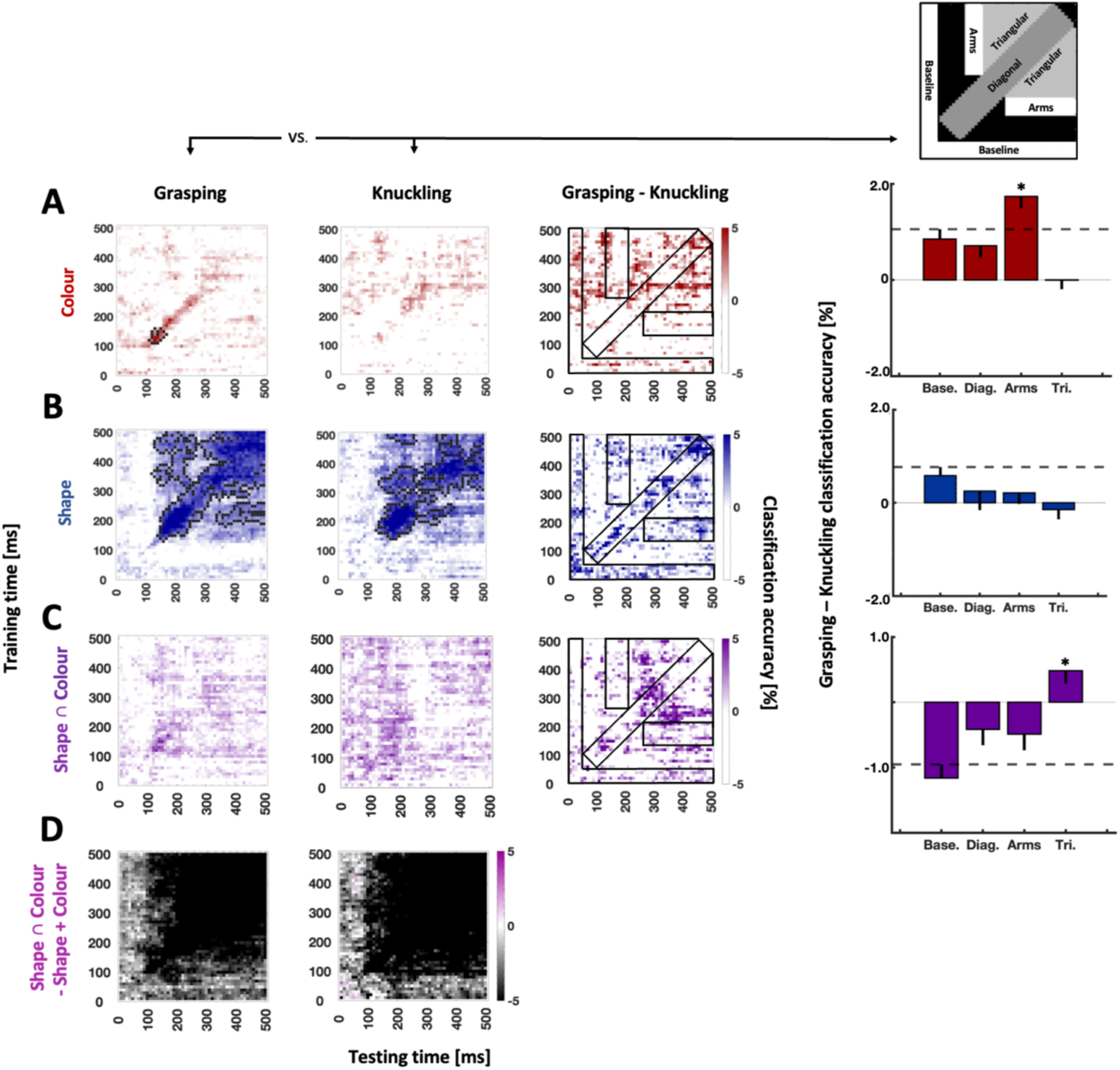
Temporal generalization of Colour (A), Shape (B), Shape ∩ Colour (C), and difference plots of Shape ∩ Colour vs. additive models of Shape and Colour (D) aligned to Preview onset. Left side: Time-by-time plots for grasping (first plot), knuckling (second plot), and the difference of the grasping and knuckling plots (third plot). Black pixels in the time-by-time plots in columns 1 and 2 mark the contours of significant clusters (cluster-based sign-permutation test with cluster-defining and cluster-size thresholds of p < 0.01). Top right graph: ROIs as defined based on Lee et al., 2024. The same ROIs are superimposed as black outlined rectangles and L-shaped areas onto the plots in the third column. Right side: Bar graphs depict observed differences between grasping and knuckling for the ROIs. Base: baseline ROI, Diag: diagonal ROI, Arms: “arms”-shaped ROI, Tri: triangular ROI. Error bars represent bootstrapped one-tailed confidence intervals (10,000 iterations, one-tailed). Dashed lines visualize the upper boundary of the confidence interval for the baseline ROI. Asterisks represent significant results relative to baseline ROI.

Grasping Shapes analyses revealed significance relative to chance both off and along the diagonal of the classification matrix (Fig. 3B, first graph). Likewise, Knuckling Shapes representations were significant both along the diagonal and in non-continuous regions off-diagonal (Fig. 3B, second graph). Comparing Grasping and Knuckling Shapes data with one another (Fig. 3B, third graph) we observed a non-significant trend for greater Grasping than Knuckling accuracy within the arms-shaped ROI. However, the trend remained below the threshold defined by the baseline ROI, and no other ROIs was significant either (Fig. 3B, fourth graph).

For Shape ∩ Colour representations we observed significantly greater accuracy for grasping than knuckling within the triangular-shaped ROIs (bar graph in Fig. 3C). However, this difference does not seem to amount to evidence for a truly integrated representation because, as shown in Fig. 4D, at almost all times Shape ∩ Colour representations were outperformed by additive models (i.e., nearly all difference values were negative).

**Figure 4.**
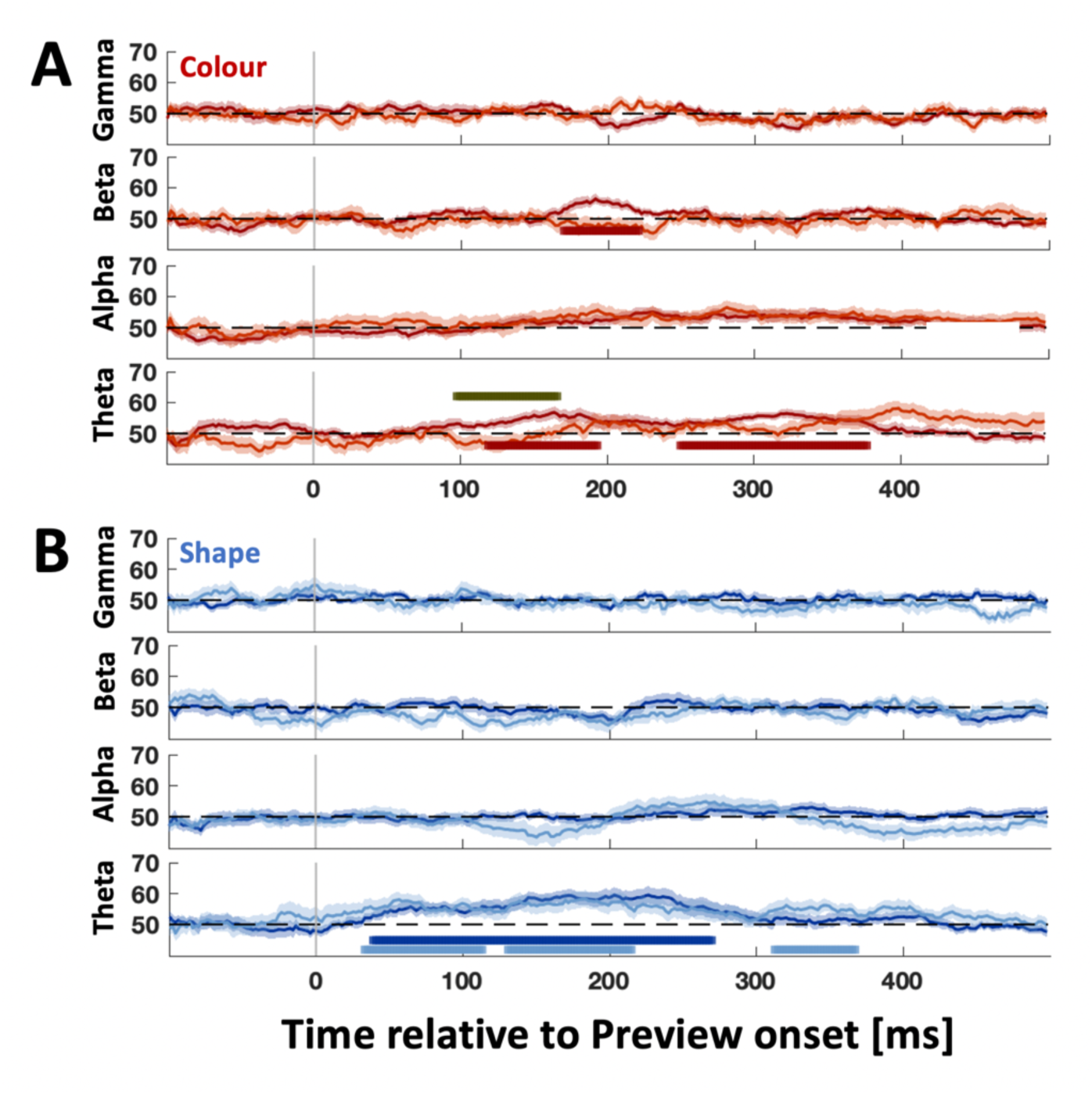
Time frequency analysis of Colour and Shape representations aligned to Preview. Time-resolved plots of Shape and Colour classification for time frequency data within separate frequency bands for Colour (A), and Shape (B). Darker curves: grasping data, lighter curves: knuckling data. Horizontal coloured bars represent above-chance classification accuracies (cluster-corrected t-test, one-tailed; p<0.05). Grey horizontal bars denote significantly better classification for grasping compared to knuckling (cluster-corrected t-test, one-tailed; p<0.05).

We also performed analyses for Grasp Orientation (refer to supplementary, S1). Due to the unbalanced nature of action conditions, the equivalent was not available for Knuckling. We see a robust sustained representation for Grasp Orientation starting as early as 40 ms to the end of the action planning period.

### Investigate Potential Sources for Grasp-Specific Reactivation Using Time Frequency Analyses: Are Cognitive Mechanisms Implicated?

To further investigate the potential oscillatory sources of our two main grasp-relevant variables, Shape and Colour, we conducted classification for separate frequency bands. In terms of time-resolved analysis of Colour (Fig. 4A), we found significance in the beta band for Grasping Colour from 170– 220 ms (maximum at 190 ms). Theta band yielded significant Grasping Colour representations from 120 – 195 ms and from 250 – 375 ms (maximum at 160 ms), with Knuckling Colour representations remaining non-significant. However, the difference between Grasping and Knuckling Colour was significant between 95 – 165 ms. There were no significant representations for alpha or gamma.

For Shape (Fig. 4B), Grasping Shape representations were significant for theta between 40 – 270 ms (maximum at 200 ms), whereas Knuckling Shape was significant from 30 – 220 ms except for an interruption from 110 – 130 ms (maximum at 180 ms), as well as significant from 310 – 370 ms. Grasping and Knuckling Shape curves were statistically not different from one another. No other frequency band yielded significant shape effects.

Next, given the significant time frequency classification results from Fig. 4 as criterion to also analyze their temporal dynamics, we performed temporal generalization analysis for Colour within theta and beta band, and for Shape within theta band to look for differences between Grasping and Knuckling.

For Colour we observed that Grasping vs. Knuckling differences within beta band produced better classification along the diagonal and the “arms”-shaped ROI, consistent with reactivation of representations (Fig. 5A). In addition, classifying within theta band yielded a significant effect along the diagonal and in the “triangular” regions of interest, consistent with sustained representations (Fig. 5B, second plot). Theta band produced no significant effect for temporal boundaries of the “arms”-shaped ROI as defined by our previous study (Lee et al., 2024). However, for exploratory reasons we also inspected an “arms”-shaped ROI for an earlier time (90 – 160 ms) given the grasp-specific difference observed for the time-resolved analysis (Fig. 4A, theta band). For this post-hoc defined ROI we did observe significantly better classification for Grasping Colour compared to Knuckling Colour (Fig. 5B, third plot). Further testing is required to confirm whether the difference reflects earlier re-activation of grasp-specific Colour representations within theta band. Lastly, Grasping Shapes vs. Knuckling Shapes within theta band yielded significantly better classification along the diagonal and “arms”-shaped ROIs, consistent with what would be expected of a reactivation pattern (Fig. 5C). For a breakdown of individual decoding accuracy differences between Grasping and Knuckling for “arms-shaped” ROIs refer to S2.

**Figure 5.**
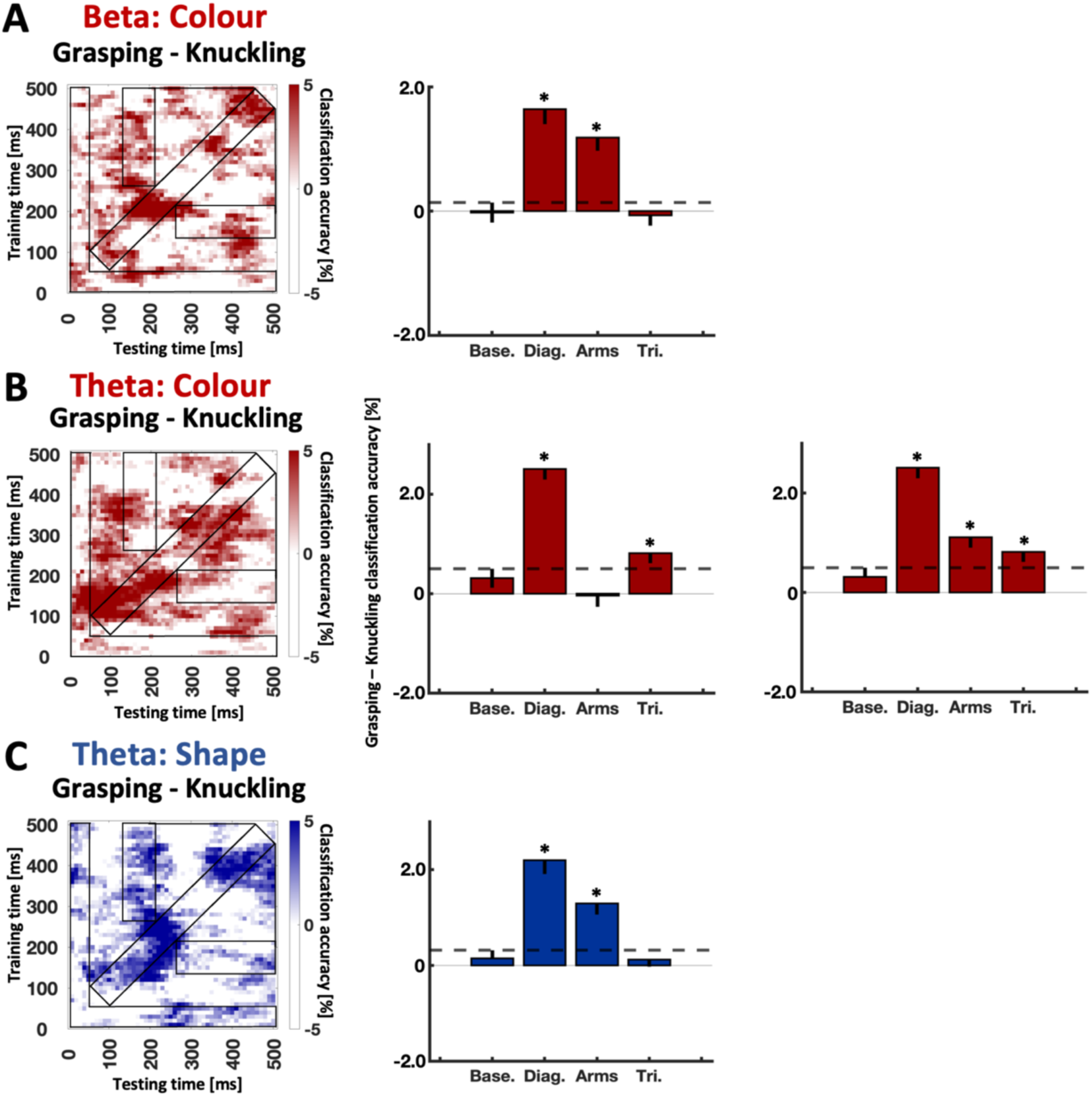
Temporal generalization of time frequency data for Grasping vs. Knuckling differences for Colour and Shape representations aligned to Preview. (A) Colour classification for beta, (B) Colour classification for theta and (C) Shape classification for theta. The outlines of the ROIs used to calculate classification differences between grasping and knuckling have been superimposed in black. Bar graphs depict differences between grasping and knuckling for the pre-defined ROIs based on Fig. 3. Rightmost bar graph depicts differences between grasping and knuckling for theta colour classification when “arms”-shaped ROIs are shifted 50 ms earlier (90 – 160 ms). Base: baseline ROI, Diag: diagonal ROI, Arms: “arms”-shaped ROI, Tri: Triangular ROI. Error bars represent bootstrapped confidence intervals (10,000 iterations, one-tailed). Horizontal dashed lines visualize significance level (i.e., the upper boundary of the confidence interval for the baseline ROI). Asterisks represent significant results relative to baseline ROI.

## Discussion

In the present study we tested how the intention to grasp an object influences cortical representations of arbitrary weight cues that are transiently stored in memory. Human participants first learned that red and blue coloured objects were made from solid and hollow blocks of plastic, respectively, or vice versa and then grasped the objects while we used EEG to record their brain activity. Training classifiers on the data, we found that representations of object colour generalized in time more strongly when participants planned to grasp the objects compared to when they intended to touch them with their knuckle.

### Cognitive Mechanisms Implicated in Reactivated Colour Representations during Grasp Planning

In detail, the task-specific difference arose within “arms-shaped” ROIs of the temporal generalization plot (Lee et al., 2024), a pattern that is indicative of earlier representations being reactivated at later times (King & Dehaene, 2014). Furthermore, we found the same reactivation pattern within beta band. Our results suggest that the colours of the objects were represented as action-specific cues in working memory in the context of motor planning extracting information about the weight of objects.

The current findings differ from previous results where representations of object material (and therefore possibly representations of weight information) occurred during similar times but were not grasp-specific (Lee et al., 2024). In the previous study, weight cues came from materials that are commonly used in daily life with overlearned material-weight contingencies. Therefore, weight information was likely recalled automatically when objects became visible, no matter what participants intended to do. In contrast, the colour-weight cues in the present study were based on newly stored contingencies. Therefore, contingencies would not have been automatically recalled – or at least such recall would have been much less probable. Instead, recall should have happened mostly, if not exclusively, when participants viewed the objects with the intention to grasp them. Consistent with this reasoning we observed reactivation of colour representations within ERPs when participants were planning to grasp objects.

What is more, the intention-triggered memory query about the respective colour-weight contingency should then have been represented in working memory. Maintaining such working memory content and top-down influences of working memory on sensory inputs have been shown to be prominently implemented within beta-band oscillations (e.g., Engel & Fries, 2010; Fiebelkorn & Kastner, 2019; Lundqvist et al., 2011; Richter et al., 2018). Indeed, here we found that classifiers trained on beta-band revealed the same grasp-specific reactivation pattern of colour as found for ERPs. In sum, this suggests that action intention triggering the recall of the colour-weight contingency resulted in a working-memory guided reactivation of neural processing of colour after 270 ms that recapitulated earlier colour processing from 140 – 210 ms, presumably to convert colour cue into weight information. This could be consistent with observations from delayed match-to-sample tasks which have found that beta modulations are correlated with successful discrimination and found later in the delay (Spitzer et al., 2010; Spitzer & Haegens, 2017; Wimmer et al., 2016) when working memory information might be reactivated for the impending comparison task. An alternate explanation might be that information contained in beta is instead linked to the maintenance of the sensorimotor set (Androulidakis et al. 2006; Pogosayan et al., 2009; Swann et al., 2009), or reflects anticipatory motor processes (Donner et al., 2009; Schoffelen et al., 2005), perhaps related to the expected sensory inputs of the colour cue. However, we will note that our previous work using a similar paradigm except with objects made of common materials did not yield grasp-specific material representations until the time of object load (Lee et al., 2024). Future work will have to disentangle the extent these processes play for representing newly learned weight associations in beta band.

Preceding the reactivation within beta band, a grasp-specific colour representations also emerged within theta band from 95 to 165 ms after object onset. The effect could mark an early onset of weight computations. But more plausible is that it reflects a shift of object- or feature-based attention to the object and its colour (e.g., Harris et al., 2017; Liebe et al., 2012). We found that these theta-band representations were significant along the diagonal of the temporal generalization plot suggesting that the representations resulted from a chain of neural generators representing (King & Dehaene, 2014) whereas our predefined “arms-shaped” ROIs (Lee et al., 2024) returned no evidence for grasp-specific reactivation. That said, it is reasonable to assume that participants shifted attention to the object’s colour before recalling the colour-weight contingency (consistent with our observation that the time-resolved grasp-specific effect within theta band in Fig. 4A occurred earlier than the temporal boundaries of the “arms-shaped” ROIs). Thus, the predefined “arms-shaped” ROIs might not have been suitable to identify an effect of reactivation. Further research is required to confirm whether the earlier ROIs as chosen post-hoc in the current study continue to yield a grasp-specific reactivation effect within theta band.

Another form of action-specific reactivation was observed for classifying shape. This was similar to our previous study (Lee et al., 2024) although in the present study reactivation occurred within theta band rather than for ERPs. This disparity might indicate that here reactivation of shape was different or perhaps weaker due to greater cognitive demands of the colour-weight contingency. Alternatively, the greater attentional demands of this experiment (i.e. keeping in mind the meaning of colours for an action planning phase that was twice as long as previous experiment) could have contributed to the diminished reactivation effect seen in ERPs that manifested instead in theta band. For example, theta is hypothesized to mediate the enhancement of target representation during encoding and retention, especially when distractors are also involved (de Vries et al., 2019; Magaosso & Borra, 2024; Riddle et al., 2020). However, once again, further research is required to examine the influence of these processes on the pattern of reactivation observed.

By contrast, the present study is largely consistent with our previous one (Lee et al., 2024) in that we found little evidence for integrated representations of object features shape and surface properties (colour vs. material), consistent with our view that computations of shape (to program movement trajectories) and weight (to program object lifting) are separate from one another (Lee et al., 2024).

In conclusion, the present study offers novel insights into interactions of cognitive and action planning processes. We show that the processing of arbitrary weight cues for later grip force computations occurs in a grasp-specific manner, arguably guided by top-down signals from working memory. These findings are consistent with a model that grasp-specific computations occur upon the computational demand of the respective action phase (Lee et al., 2024; Guo & Niemeier, 2024). The fact that reactivation of visual information occurs for multiple object features (shape, colour as well as material) and at different times (object preview/ movement planning vs. load phase, see Lee et al., 2024) seems to indicate that reactivation of earlier neural processes is a general signature of action-guided enhanced visual processing.

---

**S1.**
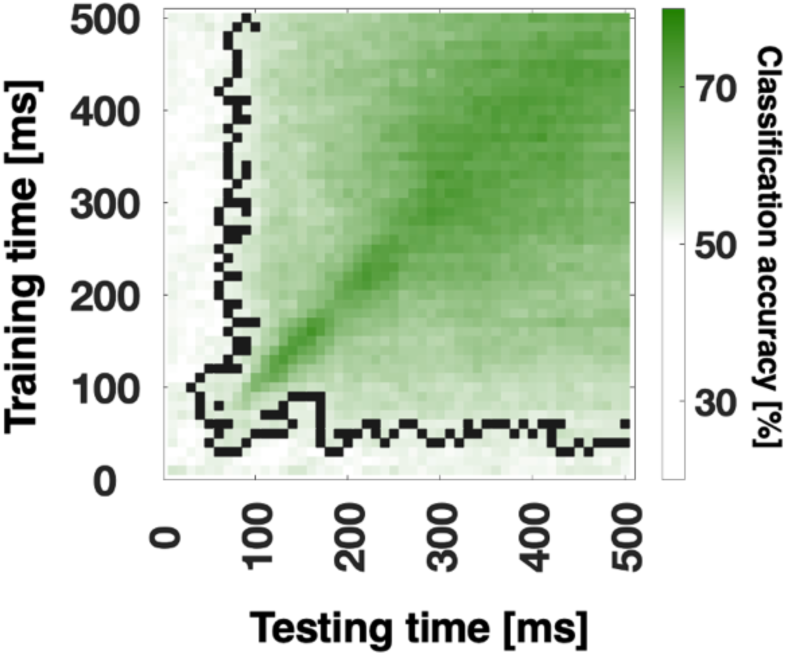
Temporal generalization of Grasp Orientation. Black pixels in the time-by-time plots in columns 1 and 2 mark the contours of significant clusters (cluster-based sign-permutation test with cluster-defining and cluster-size thresholds of p < 0.01).

**S2.**
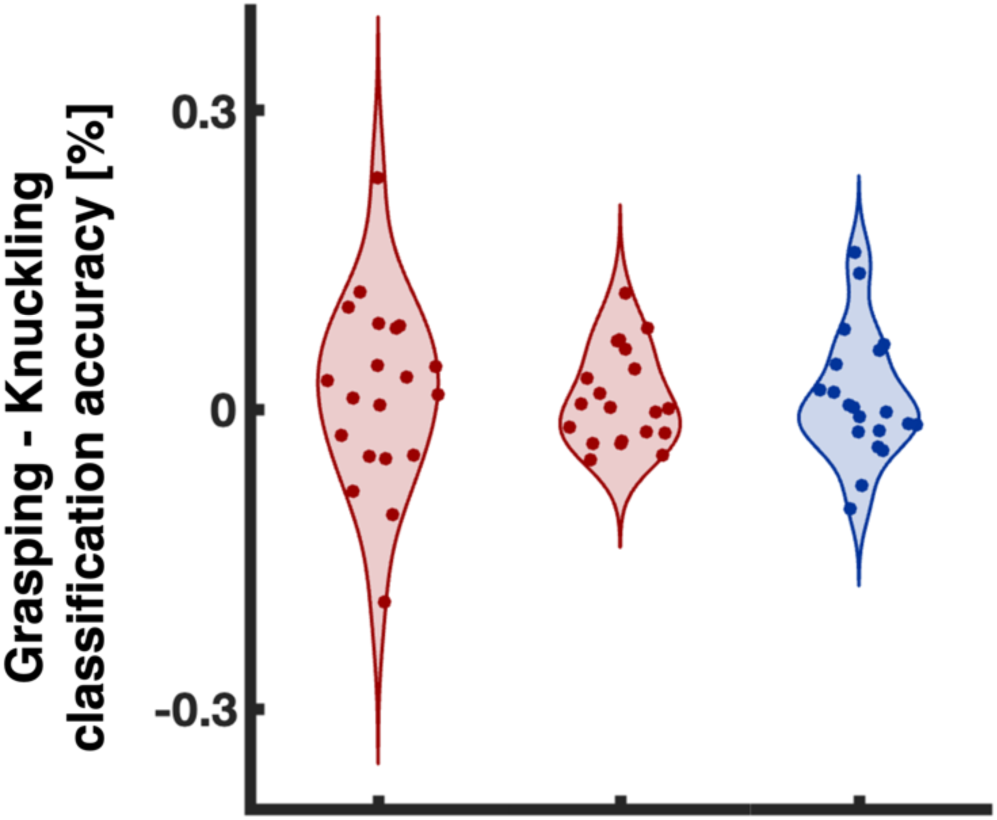
Participant difference scores between Grasping and Knuckling classification accuracies for “arms”-shaped ROIs for (from left to right): Colour classifier for ERP data, Colour classifier for beta band data, and Shape classifier for theta band data.

## Notes

**Conflict of interest statement:** The authors declare no competing financial interests.

### Competing Interest Statement

The authors have declared no competing interest.

### Summary of Updates

Title changed; Introduction updated to clarify rationale for investigating beta and theta bands; Figure 1 revised; clarification to temporal generalization Methods; Discussion updated with other possible interpretations of beta reactivation; Supplemental analyses included for grasp orientation and individual-level effects for grasp-specific reactivation.

## References

Ameli M, Dafotakis M, Fink GR, Nowak DA. Predictive force programming in the grip-lift task: the role of memory links between arbitrary cues and object weight. Neuropsychologia. 2008:46(9):2383–2388. 10.1016/j.neuropsychologia.2008.03.011.

Androulidakis AG, Doyle LM, Gilbertson TP, Brown P. Corrective movements in response to displacements in visual feedback are more effective during periods of 13–35 Hz oscillatory synchrony in the human corticospinal system. Eur J Neurosci. 2006:24:3299–3304. 10.1111/j.1460-9568.2006.05201.x

Andersen RA, Cui H. Intention, action planning, and decision making in parietal-frontal circuits. Neuron. 2009:63(5):568–583. 10.1016/j.neuron.2009.08.028.

Baldauf D, Deubel, H. Attentional selection of multiple goal positions before rapid hand movement sequences: an event-related potential study. J Cogn Neurosci. 2009:21(1), 18–29. 10.1162/jocn.2008.21021.

Baldauf D, Wolf M, Deubel, H. Deployment of visual attention before sequences of goal-directed hand movements. Vis Res. 2006:46(26), 4355–4374. 10.1016/j.visres.2006.08.021.

Batuwita R, Palade V. Class imbalance learning methods for support vector machines In: He H, Ma Y, editors. Imbalanced learning: foundations, algorithms, and applications. New York (NY): Wiley-IEEE; 2013. p. 1-15.

Baugh LA, Yak A, Johansson RS, Flanagan JR. Representing multiple object weights: competing priors and sensorimotor memories. J Neurophysiol. 2016:116(4):1615–1625. 10.1152/jn.00282.2016.

Bekkering, H, Neggers, SFW. Visual search is modulated by action intentions. Psychol Sci. 2002:13(4): 370–374. 10.1111/j.0956-7976.2002.00466.x.

Brouwer, AM, Franz, VH, Gegenfurtner, KR. Differences in fixations between grasping and viewing objects. J Vis. 2009:9(1):18. 10.1167/9.1.18.

Chaumon M, Bishop DVM, Busch NA. A practical guide to the selection of independent components of the electroencephalogram for artifact correction. J Neurosci Methods. 2015:250:47–63. 10.1016/j.jneumeth.2015.02.025.

Craighero L, Fadiga L, Rizzolatti G, & Umiltà C. Action for perception: a motor–visual attentional effect. J of Exp Psychol Hum Percept Perform. 1999:25:1673–1692. 10.1037/0096-1523.25.6.1673.

Cui H. From intention to action: hierarchical sensorimotor transformation in the posterior parietal cortex. eNeuro. 2014:1(1):ENEURO.0017-14.2014. 10.1523/ENEURO.0017-14.2014.

de Vries IEJ, Savran E, van Driel J, Olivers CNL. Oscillatory mechanisms of preparing for visual distraction. J Cogn Neurosci. 2019:31:1873–1894. 10.1162/jocn_a_01460

Delorme A, Makeig S. EEGLAB: an open source toolbox for analysis of single-trial EEG dynamics including independent component analysis. J Neursci Methods. 2004:134:9–21. 10.1016/j.jneumeth.2003.10.009.

Donner TH, Siegel M, Fries P, Engel AK. Buildup of choice predictive activity in human motor cortex during perceptual decision making. Curr Biol. 2009:19:1581–1585. 10.1016/j.cub.2009.07.066

Drisdelle BL, Aubin S, Jolicoeur P. Dealing with ocular artifacts on lateralized ERPs in studies of visual-spatial attention and memory: ICA correction versus epoch rejection. Psychophysiol. 2017: 54(1):83–99. 10.1111/psyp.12675.

Engel AK, Fries P. Beta-band oscillations--signalling the status quo? Curr Opin Neurobiol. 2010:20(2):156–165. 10.1016/j.conb.2010.02.015.

Fagioli, S, Hommel, B, Schubotz, RI. Intentional control of attention: action planning primes action-related stimulus dimensions. Psychol Res. 2007:71: 22–29. 10.1007/s00426-005-0033-3.

Fellrath J, Mottaz A, Schnider A, Guggisberg AG, Ptak R. Theta-band functional connectivity in the dorsal fronto-parietal network predicts goal-directed attention. Neuropsychologia. 2016:92, 20–30. 10.1016/j.neuropsychologia.2016.07.012

Fiebelkorn IC, Kastner S. A rhythmic theory of attention. Trends Cogn Sci. 2019:23(2):87–101. 10.1016/j.tics.2018.11.009.

Gallivan JP, Cant JS, Goodale MA, Flanagan JR. Representation of object weight in human ventral visual cortex. Curr Biol. 2024:24:1866–1873. 10.1016/j.cub.2014.06.046.

Gallivan, JP, Chapman, CS, Gale, DJ, Flanagan, JR, & Culham, JC. Selective modulation of early visual cortical activity by movement intention. Cereb Cortex. 2019:29(11):4662–4678. 10.1093/cercor/bhy345.

Guillery E, Mouraux A, Thonnard JL, Legrain V. Mind your grip: even usual dexterous manipulation requires high level cognition. Front Behav Neurosci. 2017:11:220. 10.3389/fnbeh.2017.00220.

Grootswagers T, Wardle SG, Carlson TA. Decoding dynamic brain patterns from evoked responses: a tutorial on multivariate pattern analysis applied to time series neuroimaging data. J Cog Neurosci. 2017:39:677–697. 10.1162/jocn_a_01068.

Guggenmos M, Sterzer P, Cichy RM. Multivariate pattern analysis for MEG: a comparison of dissimilarity measures. Neuroimage. 2018:173:434–447. 10.1016/j.neuroimage.2018.02.044.

Guo LL, Nestor A, Nemrodov D, Frost A, Niemeier M. Multivariate analysis of electrophysiological signals reveals the temporal properties of visuomotor computations for precision grips. J Neurosci. 2019:39(48):9585–9597. 10.1523/JNEUROSCI.0914-19.2019.

Guo LL, Niemeier M. Phase-dependent visual and sensorimotor integration of features for grasp computations before and after effector specification. J Neurosci. 2024:44(33):e2208232024. 10.1523/JNEUROSCI.2208-23.2024.

Guo LL, Oghli YS, Frost A, Niemeier M. Multivariate analysis of electrophysiological signals reveals the time course of precision grasps programs: evidence for nonhierarchical evolution of grasp control. J Neurosci. 2021:41(44):9210–9222. https://10.1523/JNEUROSCI.0992-21.2021.

Harris AM, Dux PE, Jones CN, Mattingley JB. Distinct roles of theta and alpha oscillations in the involuntary capture of goal-directed attention. Neuroimage. 2017:152:171–183. 10.1016/j.neuroimage.2017.03.008.

Hsieh L-T, Ranganath C. Frontal midline theta oscillations during working memory maintenance and episodic encoding and retrieval. NeuroImage. 2014:85: 721–729. 10.1016/j.neuroimage.2013.08.003.

Jung TP, Makeig S, Humphries C, Lee TW, McKeown MJ, Iragui V, Sejnowski TJ. Removing electroencephalographic artifacts by blind source separation. Psychophysiol. 2000:37:163–178. 10.1111/1469-8986.3720163.

King JR, Dehaene, S. Characterizing the dynamics of mental representations: the temporal generalization method. Trends Cogn Sci. 2014:18:203–210. 10.1016/j.tics.2014.01.002.

Klein LK, Maiello G, Stubbs K, Proklova D, Chen J, Paulun VC, Culham JC, Fleming RW. Distinct neural components of visually guided grasping during planning and execution. J Neurosci. 2023:43:8504–8514. 10.1523/JNEUROSCI.0335-23.2023.

Lee N, Guo LL, Nestor A, Niemeier M. Computation on demand: action-specific representations of visual task features arise during distinct movement phases. J Neurosci. 2024:44(29):e2100232024. 10.1523/JNEUROSCI.2100-23.2024.

Li Y, Randerath J, Bauer H, Marquardt C, Goldenberg G, Hermsdörfer J. Object properties and cognitive load in the formation of associative memory during precision lifting. Behav Brain Res. 2009:196(1):123–130. 10.1016/j.bbr.2008.07.031.

Liebe S, Hoerzer GM, Logothetis NK, Rainer G. Theta coupling between V4 and prefrontal cortex predicts visual short term memory performance. Nat Neurosci. 2012:15:456–462. 10.1038/nn.3038.

Lopez-Calderon J, Luck SJ. ERPLAB: an open-source toolbox for the analysis of event-related potentials. Front Hum Neurosci. 2014:8:213. 10.3389/fnhum.2014.00213.

Lundqvist M, Herman P, Lansner A. Theta and gamma power increases and alpha/beta power decreases with memory load in an attractor network model. J Cogn Neurosci. 2011:23(10):3008–3020. 10.1162/jocn_a_00029.

Magosso E, Borra D. The strength of anticipated distractors shapes EEG alpha and theta oscillations in a working memory task. NeuroImage. 2024:300:120835. 10.1016/j.neuroimage.2024.120835

Maslow, AH. The psychology of science: a reconnaissance. 1st ed. New York (NY): Harper & Row; 1966.

Miller EK, Lundqvist M, Bastos AM. Working memory 2.0. Neuron. 2018:100(2):463–475. 10.1016/j.neuron.2018.09.023.

Nemrodov D, Niemeier M, Patel A, Nestor A. The dynamics of facial identity processing: an EEG-based image reconstruction study. J Vis. 2017:17:1262. 10.1167/17.10.1262.

Nemrodov D, Niemeier M, Patel A, Nestor A. The neural dynamics of facial identity processing: insights from EEG-based pattern analysis and image reconstruction. eNeuro. 2018:5: ENEURO.0358-17.2018. 10.1523/eneuro.0358-17.2018.

Nichols TE, Holmes AP. Nonparametric permutation tests for functional neuroimaging: a primer with examples. Hum Brain Mapp. 2002:15:1–25. 10.1002/hbm.1058.

Oldfield RC. The assessment and analysis of handedness: the Edinburgh inventory. Neuropsychologia. 1971:9(1):97–113. 10.1016/0028-3932(71)90067-4.

Pogosyan A, Gaynor LD, Eusebio A, Brown P. Boosting cortical activity at beta-band frequencies slows movement in humans. Curr Biol. 2009:19:1637–1641. 10.1016/j.cub.2009.07.074

Richter CG, Coppola R, Bressler SL. Top-down beta oscillatory signaling conveys behavioral context in early visual cortex. Sci Rep. 2018:8(1):6991. 10.1038/s41598-018-25267-1.

Riddle J, Scimeca JM, Cellier D, Dhanani S, D’Esposito M. Causal evidence for a role of theta and alpha oscillations in the control of working memory. Curr. Biol. 2020:30: 1748–1754.e4. 10.1016/j.cub.2020.02.065

Sauseng, P, Griesmayr, B, Freunberger, R, & Klimesch, W. Control mechanisms in working memory: A possible function of EEG theta oscillations. Neurosci Biobehav Rev. 2010:34(7), 1015–1022. 10.1016/j.neubiorev.2009.12.006

Schiegg, A, Deubel, H, Schneider, W. Attentional selection during preparation of prehension movements. Vis Cogn. 2003:10(4):409–431. 10.1080/13506280244000140.

Schoffelen JM, Oostenveld R, Fries P. Neuronal coherence as a mechanism of effective corticospinal interaction. Science. 2005:308:111–113. 10.1126/science.1107027

Shin, YK, Proctor, RW, Capaldi, EJ. “A review of contemporary ideomotor theory”: correction to Shin et al. (2010). Psychol. Bull. 2010:136(6):974. 10.1037/a0021628.

Spitzer B, Gloel M, Schmidt TT, Blankenburg F. Working memory coding of analog stimulus properties in the human prefrontal cortex. Cereb Cortex. 2013:24(8), 2229–2236. 10.1093/cercor/bht084

Spitzer B, Haegens S. Beyond the status quo: A role for beta oscillations in endogenous content (re)activation. eNeuro, 2017:4(4), ENEURO.0170-17.2017. 10.1523/eneuro.0170-17.2017

Spitzer B, Wacker E, Blankenburg F. Oscillatory correlates of vibrotactile frequency processing in human working memory. J Neurosci. 2010:30(12), 4496–4502. 10.1523/jneurosci.6041-09.2010

Swann N, Tandon N, Canolty R, Ellmore TM, McEvoy LK, Dreyer S, DiSano M, Aron AR. Intracranial EEG reveals a time- and frequency-specific role for the right inferior frontal gyrus and primary motor cortex in stopping initiated responses. J Neurosci. 2009:29:12675–12685. 10.1523/JNEUROSCI.3359-09.2009

Tesche CD, Karhu J. Theta oscillations index human hippocampal activation during a working memory task. Proc. Natl. Acad. Sci. U.S.A. 2000:97(2), 919–924. 10.1073/pnas.97.2.919

Wimmer K, Ramon M, Pasternak T, Compte A. Transitions between multiband oscillatory patterns characterize memory-guided perceptual decisions in prefrontal circuits. J Neurosci. 2016:36:489–505. 10.1523/JNEUROSCI.3678-15.2016 pmid:26758840

Zhang Z, Cesanek E, Ingram JN, Flanagan JR, Wolpert DM. Object weight can be rapidly predicted, with low cognitive load, by exploiting learned associations between the weights and locations of objects. J Neurophysiol. 2023:129(2):285–297. 10.1152/jn.00414.2022.

